# HOIL1 regulates group 3 innate lymphoid cell numbers in the colon and protects against systemic dissemination, colonic ulceration, and lethality from *Citrobacter rodentium* infection

**DOI:** 10.1101/2023.05.22.541746

**Authors:** Victoria L. Hartley, Arwa M. Qaqish, Matthew J. Wood, Brian T. Studnicka, Kazuhiro Iwai, Ta-Chiang Liu, Donna A. MacDuff

**Author notes:** Address correspondence to Donna A. MacDuff.

## Abstract

HOIL1-deficient patients experience chronic intestinal inflammation and diarrhea as well as increased susceptibility to certain bacterial infections. HOIL1 is a component of the linear ubiquitin chain assembly complex (LUBAC) that regulates immune signaling pathways including NF-κB-activating pathways. We have shown previously that HOIL1 is essential for survival following *Citrobacter rodentium* gastrointestinal infection of mice, but the mechanism of protection by HOIL1 was not examined. *C. rodentium* is a murine model for human attaching and effacing (A/E) pathogens, enteropathogenic and enterohemorrhagic *Escherichia coli,* that cause diarrhea and food-borne illnesses, and lead to severe disease in children and immunocompromised individuals. In this study, we found that *C. rodentium* infection caused severe colitis and dissemination of *C. rodentium* to systemic organs in *Hoil1^-/-^*mice. HOIL1 was important in radiation-resistant cells and in the innate immune response to limit early replication of *C. rodentium* in the intestine, and to modulate induction of inflammatory cytokines. Using cell type-specific knock-out mice, we found that HOIL1 was dispensable in intestinal epithelial cells (IEC), but was required in CD11c- and lysozyme 2-expressing myeloid cells to prevent weight loss and systemic dissemination of *C. rodentium*. While HOIL1-deficiency did not affect populations of neutrophils or macrophages, dendritic cells and group 3 innate lymphoid cell (ILC3) numbers were reduced, resulting in a defect in IL-22 induction during *C. rodentium* infection. Understanding the role HOIL1 plays in limiting the pathogenesis of A/E lesion-forming bacteria will provide further insights into the innate immune response to gastrointestinal pathogens and inflammatory disorders.

## Introduction

Enterohemorrhagic *Escherichia coli* (EHEC) and enteropathogenic *E. coli* (EPEC) infections are a leading cause of bacterial diarrheal disease and lethality in children in the developing world (1, 2). In developed countries, EPEC and EHEC are common causative agents of foodborne illnesses (3). While disease symptoms are typically mild in immunocompetent adults, infection can lead to severe morbidity and mortality in infants and immunocompromised individuals (3–5). Since EPEC and EHEC do not easily infect mice, their murine counterpart, *Citrobacter rodentium,* has been used extensively to model and investigate these pathogens *in vivo* (6, 7). EPEC, EHEC, and *C. rodentium* infections are characterized by the formation of attaching and effacing (A/E) lesions on intestinal epithelial cells (8), wherein bacteria attach directly to intestinal epithelium, efface the microvilli, and form pedestal-like structures on the epithelial cells beneath the adherent bacteria. This process is mediated by a bacterial type III secretion system that injects effector proteins into the intestinal epithelial cells, altering their metabolism, actin polymerization (9) and immune functions (7, 10, 11). *C. rodentium-*induced colitis is also used as an experimental animal model for inflammatory bowel diseases such as Crohn’s disease and ulcerative colitis (12).

Following peroral inoculation of C57BL/6 mice, *C. rodentium* initially colonizes the cecum followed by the colon within two to three days (13). *C. rodentium* infection induces colonic colitis and hyperplasia, characterized by infiltration of immune cells, excessive proliferation of intestinal epithelial cells (IEC) and deepening of colonic crypts. Bacterial burdens are mostly contained within the GI tract, peaking around six to ten days post-infection, and are cleared within three weeks (13). Upon infection, pattern recognition receptors such as Toll-like receptors (TLRs) on IEC, dendritic cells and macrophages in the intestine detect the presence of pathogenic bacteria and initiate innate immune responses. Production of cytokines such as IL-23 and IL-1β by dendritic cells and macrophages leads to the production of IL-22 by group 3 innate lymphoid cells (ILC3) and T helper type 22 (Th22) cells (14–19). IL-22 is crucial for the early mucosal immune response to control *C. rodentium*, as it promotes barrier defense through IEC proliferation and the production of antimicrobial peptides (17, 20–22). The adaptive immune response, particularly the production of antibodies against *C. rodentium,* is essential for clearance of the pathogen and survival following infection (23).

We have shown previously that heme-oxidized IRP2 ubiquitin ligase-1 (HOIL1, official gene name *Rbck1*), a component of the Linear Ubiquitin Chain Assembly Complex (LUBAC), is essential for survival during *C. rodentium* infection (24). However, the mechanism by which HOIL1 protects against *C. rodentium*-induced pathogenesis is unknown. LUBAC is a trimeric complex composed of HOIL1, HOIP and SHARPIN. The catalytic subunit, HOIP, is the only E3 ubiquitin ligase known to generate linear (methionine-1 linked) ubiquitin chains (25–27). LUBAC has been implicated in the regulation of type 1 inflammatory cytokines downstream of multiple innate immune receptors that lead to the activation of canonical NF-κB, such as TNFR1, IL-1- and IL-17-family receptors, TLRs, and CD40, as well as in the modulation of programmed cell death pathways (24, 26–36). HOIL1 also contains a functional E3 ubiquitin ligase domain (37, 38). However, the range of substrates ubiquitinated by HOIL1 and their downstream functions are not well understood (37–39). In humans, HOIL1 deficiency is associated with a lethal disorder characterized by increased susceptibility to pyogenic bacterial infection, chronic autoinflammation, muscular amylopectinosis and IBD-like symptoms (40–42). Since full deletion of *Hoil1* leads to embryonic lethality in mice (43, 44), our lab uses a mouse strain that express a truncated version of HOIL1 lacking the C-terminal RING-between-RING domain to study the physiological functions of HOIL1 *in vivo* (24, 26, 43). Expression of HOIP and SHARPIN is also reduced in these mice. While these “HOIL1-mutant” mice (*Hoil1^-/-^* herein) display no overt phenotype when housed in specific pathogen-free (SPF) conditions, they are highly susceptible to certain infections including *Listeria monocytogenes, Toxoplasma gondii,* and *C. rodentium*. Additionally, *Hoil1^-/-^* mice exhibit hyperinflammation during chronic infection with *Mycobacterium tuberculosis* or murine gamma-herpesvirus 68, as well as progressive amylopectinosis in their cardiac tissue, suggesting that they are a suitable model for HOIL1-deficiency in humans (24, 40, 41). While susceptibility of *Hoil1^-/-^* mice to *Listeria* infection is associated with defective induction of type 1 inflammatory cytokines, including IL-12, TNF-α, IL-6 and interferon (IFN)-γ, the role of HOIL1 in the type 3 inflammatory response to extracellular bacterial pathogens such as to *C. rodentium* has not been elucidated.

Here, we show that HOIL1 plays an essential role in the innate immune response to limit early *C. rodentium* replication in the GI tract, as well as to prevent dissemination to systemic tissues and severe colonic ulceration. HOIL1 modulated the inflammatory cytokine response, including IL-22, during *C. rodentium* infection. Using cell type-specific knock-out mice, we show that HOIL1 plays important roles in CD11c-expressing and lysozyme-2-expressing myeloid cells, but not in Villin 1-expressing IECs, to prevent systemic dissemination and weight loss. However, HOIL1 is not required to regulate populations of these cells before or during infection. Instead, we found that HOIL1 regulates ILC populations in the colon, leading to a decrease the proportion of RORγt^+^ ILC3 in the colon at baseline, and an impaired IL-22 response during infection. These findings highlight a key role for HOIL1 in modulating innate mucosal immunity to an A/E gastrointestinal pathogen.

## Materials and Methods

### Mice

*Hoil1^-/-^* (B6.Cg-Hoil1^tm1Kiwa^), *Rag1^-/-^Hoil1^-/-^* and *Hoil1^-/-^Il4ra^ΔIEC^* mice have been described previously (24, 26, 45). Wild type control mice were either *Hoil1^+/+^* littermates or C56BL/6 mice from our colony. All phenotypes were confirmed with *Hoil1^+/+^* littermates. *Rag1^-/-^Hoil1^+/+^*littermates were used as controls for *Rag1^-/-^Hoil1^-/-^*mice. Heterozygous B6NCrl;B6N-A^tm1Brd^Rbck1^tm1a(EUCOMM)Hmgu^ mice were acquired from INFRAFRONTIER/EMMA (European mutant mouse archive) (46, 47). Mice were crossed to FLP deleter mice to remove the targeting cassette to generate *Rbck1*^+/f^ mice, then backcrossed to B6J using speed congenics. *Rbck1*^+/f^ mice were crossed to B6.C-Tg(CMV-cre)1Cgn/J mice (Cre deleter, Jackson labs) to delete the floxed region. *Rbck1*^+/-^ mice were crossed to *Rbck1*^f/f^ and generate *Rbck1*^f/-^ mice. Mice were crossed to B6.Cg-Tg(Vil1-cre)1000Gum/J to generate *Rbck1*^f/-Vil1-cre^ mice. Mice were crossed to B6.Cg-Tg(Itgax-cre)1-1Reiz/J mice to generate *Rbck1*^f/-CD11c-cre^ mice. Mice were crossed to B6.129P2-Lyz2tm1(cre)Ifo/J to generate *Rbck1*^f/-Lyz2-cre^ mice. Mice were age- and sex-matched for individual experiments. Mice were cohoused and littermate controls were used. Both male and female mice were used, as sex differences were not observed. Mice were housed and bred at Washington University in Saint Louis or at University of Illinois Chicago in specific pathogen-free conditions in accordance with Federal and University guidelines, and protocols were approved by the Animal Studies Committee of Washington University or the Animal Care Committee of the University of Illinois Chicago.

### *In vivo* infections

Kanamycin-resistant *C. rodentium* strain DBS120 was used for this study (48). Bacteria were grown in Luria-Bertani (LB) medium containing 50 µg/ml kanamycin at 37°C. Mice between six and eight weeks-of-age were inoculated intragastrically with 2x10^9^ CFU from a log-phase culture in 100 μl 3% sodium bicarbonate in PBS. For intraperitoneal infections, mice were injected with 2x10^7^ CFU *C. rodentium* in 300ul PBS into the peritoneal cavity. Mice were monitored daily for morbidity and mortality, and mice that had lost more than 20% of their initial body weight were euthanized.

Stool samples were collected every two days and analyzed for colony forming units (CFU). Mice that were not shedding detectable amounts of *C. rodentium* by 4 dpi were excluded from the study. Upon euthanasia, organ pieces were placed in pre-weighed sterile tubes containing 1.0 mm zirconia/silica beads (BioSpec Products Inc.) on ice for analysis of CFU, or snap frozen and stored at -80°C for future RNA or protein analysis. To determine CFU, organ and stool samples were homogenized in 1 ml PBS plus 0.1% Tween 20 with a mini-beadbeater (BioSpec Products Inc.), serially diluted in PBS plus 0.1% Tween 20, and plated on LB agar containing 50 µg/ml kanamycin.

### Generation of bone marrow chimeric mice

Bone marrow transplants were performed as described previously (24). Briefly, recipient mice were exposed to 1200 rad of whole body irradiation, and injected intravenously with 10 million whole bone marrow cells from donor mice. Mice were allowed to reconstitute for 16 weeks before inoculation with *C. rodentium*. Mice were bled at 14 weeks post-irradiation to determine percent chimerism by quantitative PCR (qPCR) on genomic DNA isolated from peripheral blood as described previously (24).

### Quantitative reverse-transcriptase-PCR

Frozen tissue samples were homogenized in 1 ml TRI Reagent (Sigma) and RNA was isolated according to the manufacturer’s instructions. RNA samples were treated with Turbo DNA-free DNase (Ambion). cDNA synthesis was performed using Improm-II (Promega) and random hexamer primers. Quantitative PCR was performed using a StepOnePlus or QuantStudio 3 real-time PCR system (Applied Biosystems) with Amplitaq Gold polymerase (Applied Biosystems) and PrimeTime probe-based qPCR Assays (Integrated DNA Technologies) specific for *Cxcl1* (assay ID: Mm.PT.58.42076891), *Il23a* (assay ID: Mm.PT.58.10594618.g), *Il6* (assay ID: Mm.PT.58.10005566), *Tnfa* (assay ID: Mm.PT.58.12575861), *Il1b* (assay ID: Mm.PT.58.41616450), *Il12b* (assay ID: Mm.PT.58.12409997), *Ifng* (assay ID: Mm.Pt.58.41769240), *Il13* (assay ID: Mm.PT.58.31366752) and *Il22* (F: AGA ACG TCT TCC AGG GTG AA; R: TCC GAG GAG TCA GTG CTA A; Probe: /56-FAM/TGA GCA CCT GCT TCA TCA GGT AGC A/36-TAMSp/). Transcripts were quantitated using specific standard curves and copy numbers normalized to the reference gene ribosomal protein S29 (*Rps29*; forward primer 5′-GCA AAT ACG GGC TGA ACA TG-3′, reverse primer 5′-GTC CAA CTT AAT GAA GCC TAT GTC-3′, and probe 5’-/5HEX/CCT TCG CGT/ZEN/ ACT GCC GGA AGC/ 3IABkFQ/-3’ (Integrated DNA Technologies).

### Histological analyses

Intestinal tissue was flushed with PBS and fixed in 10% neutral-buffered formalin at 4°C for 20 hr, washed three times with 70% ethanol and incubated in 70% ethanol at 4°C for at least 24 hr. Colons and cecums were cut into strips, embedded in 2% agar for optimal crypt orientation prior to being embedded in paraffin, sectioned, and stained with H&E. Imaging was performed on a BZ-X710 microscope (Keyence). Histologic severity scores were assigned based on a pre-determined, semi-quantitative histologic severity score (PMID: 29496999). Briefly, 0: no neutrophils; 1: neutrophils in surface epithelium, lamina propria, or cryptitis; 2: crypt abscess; 3: neutrophilic inflammation present at muscularis mucosae or submucosa; 4: ulcer or transmural inflammation; 5: epithelial denudement. The percentages of longitudinal extent of involved colon were also quantified.

### Intestinal permeability analyses

150 μl of 80 mg/ml fluorescein isothiocyanate dextran (FITC-dextran, FD4, Sigma) in PBS was administered to mice by oral gavage. 4 hr after administration, mice were euthanized with CO_2_, and blood was collected by cardiac puncture, transferred to a Microtainer Serum Separator tube (BD), and processed according to the manufacturer’s instructions. FITC-dextran concentration in the serum was determined by spectrophotofluorometry with an excitation of 485 nm and emission wavelength of 528 nm and calculated using a standard curve of BSA. Plates were read on a BioTek Synergy 2 plate reader.

### Cytokine analysis

Tissues were harvested and stored as described above. For protein analysis, tissue was homogenized using a mini-beadbeater (BioSpec Products Inc.) for 1 minute in a buffer containing 20 nM Tris HCl (pH 7.5), 150 nM NaCl, and 0.05% Tween 20 with Halt protease inhibitors (ThermoFisher Scientific). 1 ml of buffer was added to the samples per 100 mg of tissue. The samples were centrifuged to remove debris and supernatant was collected for analysis. Protein concentration was determined using DC Protein Concentration Assay (BioRad). Protein levels of cytokines were determined using Milliplex MAP Kit Mouse Th17 Magnetic Bead Panel (Cat # MTH17MAG-47K) according to manufacturer instructions. The plates were read using Magpix (Luminex).

### Flow cytometry

Mice were euthanized, and colons were removed and flushed with cold PBS. Colons were opened lengthwise and cut into 1 cm pieces. Colon pieces were incubated in HBSS buffer (supplemented with 15 mM HEPES, 5mM EDTA, 1.25 mM DTT and 10% bovine calf serum) for 20 min at RT, then vortexed in PBS to remove epithelial cells. The HBSS wash step was repeated, and then tissue was vortexed in PBS three times. Tissue pieces were placed in R10 (RPMI 1640 supplemented with 10% FBS, L-glutamine, and penicillin/streptomycin) with 0.5 mg/ml collagenase VIII (Sigma) and placed in a shaking incubator at 37°C for 15 min. Samples were shaken by hand and filtered through 100 μm strainers on ice. Cells were washed with 35 ml cold R10, centrifuged at 400g for 10 min, then resuspended in FACS buffer (PBS with 2 mM EDTA and 1% FBS). 1-2 million cells were incubated with FACS buffer with 1% Fc block, 1% mouse serum, 1% rat serum, and 1% hamster serum, then stained with fluorophore conjugated antibodies against: CD45 (30-F11), Ly6C (HK1.4), Ly6G (1A8), CD103 (2E7), CX3CR1 (SA011F1), CD11c (N418), CD11b (M1/70 and M1/70), F4/80 (BM8), CCR6 (29-21.17), NK1.1 (PK136), KLRG1 (2F1), CD90.2 (30-H12), NKp46 (29A1.4). To identify ILC, cells were stained for CD3 (17A2), CD5 (53-7.3), B220 (RA3-6B2), CD19 (6D5), TCRβ (H57-597), TCRγδ (UC7-13D5) and selecting lineage negative cells. Cells were stained with Zombie NIR fixable viability dye (BioLegend) to identify live cells. Alternatively, following digestion, cells were passed through Percoll gradient to isolate lymphocytes. The cells were resuspended in 1 ml R10 and 1-2 million cells were plated in 1ml R10 with 1 μl Brefeldin A (BioLegend) for 4 hours. Cells were collected, washed in FACS buffer, and stained with surface antibodies as described above. For intracellular staining, cells were fixed and permeabilized using FoxP3 fix/perm reagents (eBioscience) following manufacturer instructions and stained with antibodies against RORγt (BD2) and IL-22 (Poly5164) or isotype control (RTK2071). Flow cytometry was performed on a CytoFLEX S (Beckman Coulter) and data were analyzed using FlowJo (TreeStar Inc.).

### Western blot analyses

Tissue samples were collected and stored as described above. Frozen samples were homogenized in 200 μl RIPA buffer (150 mM NaCl, 50 mM Tris pH 7.4, 0.1% SDS, 1% IGEPAL (Alfa Aesar), 0.5% DOC)) with Halt protease inhibitors (ThermoFisher Scientific) per 10 mg of tissue and centrifuged for 15 minutes at 4°C at 21130 g. Supernatant was collected and diluted 1:10 in RIPA buffer, then mixed with equal volume of 2x Laemmli buffer and heated to 100°C for 5 minutes. Antibodies specific for RBCK1 (Santa Cruz Biotechnology sc-393754), HOIP (VWR/Proteintech 16298-1-AP, SHARPIN (VWR/Proteintech 14626-1-AP), and actin (Sigma-Aldrich A5316) were diluted 1:1000 in 3% milk TBS-T (HOIP, SHARPIN, actin) or 5% milk TBS-T (RBCK1). Blots were imaged using a BioRad ChemiDock Imager.

### Statistical analyses

Statistical significance was determined using GraphPad Prism 9 software as described in the figure legends.

## Results

### HOIL1 restricts *C. rodentium* intestinal growth and dissemination to systemic sites

While C57BL/6 mice exhibit mild symptoms upon gastrointestinal infection with *C. rodentium*, HOIL1-deficiency confers susceptibility to infection with weight-loss beginning around six days, and lethality occurring between 10 and 15 days post-infection (dpi; (24)). To gain further insight into role of HOIL1 in protecting against *C. rodentium* pathogenesis, we measured *C. rodentium* shedding in stool and burdens in tissues over the course of infection. Elevated *C. rodentium* shedding in the stool of *Hoil1^-/-^* mice was detectable as early as 2 dpi and continued for as long as stool pellets were obtainable from the HOIL1-mutant mice (Fig. 1A). Consistently, increased *C. rodentium* CFU were detected in the cecal contents and attached to the cecal tissue by 24 hours post-infection (hpi) (Fig. 1B). The level of colonization of the distal colon was similar in both groups of mice up to 8 dpi (Fig. 1C). While *C. rodentium* CFU were rarely detected in the spleen and liver of *Hoil1*^+/+^ mice, bacteria were detected in these tissues in almost all *Hoil1*^-/-^ mice by 4 dpi (Fig. 1D-E).

**Figure 1:**
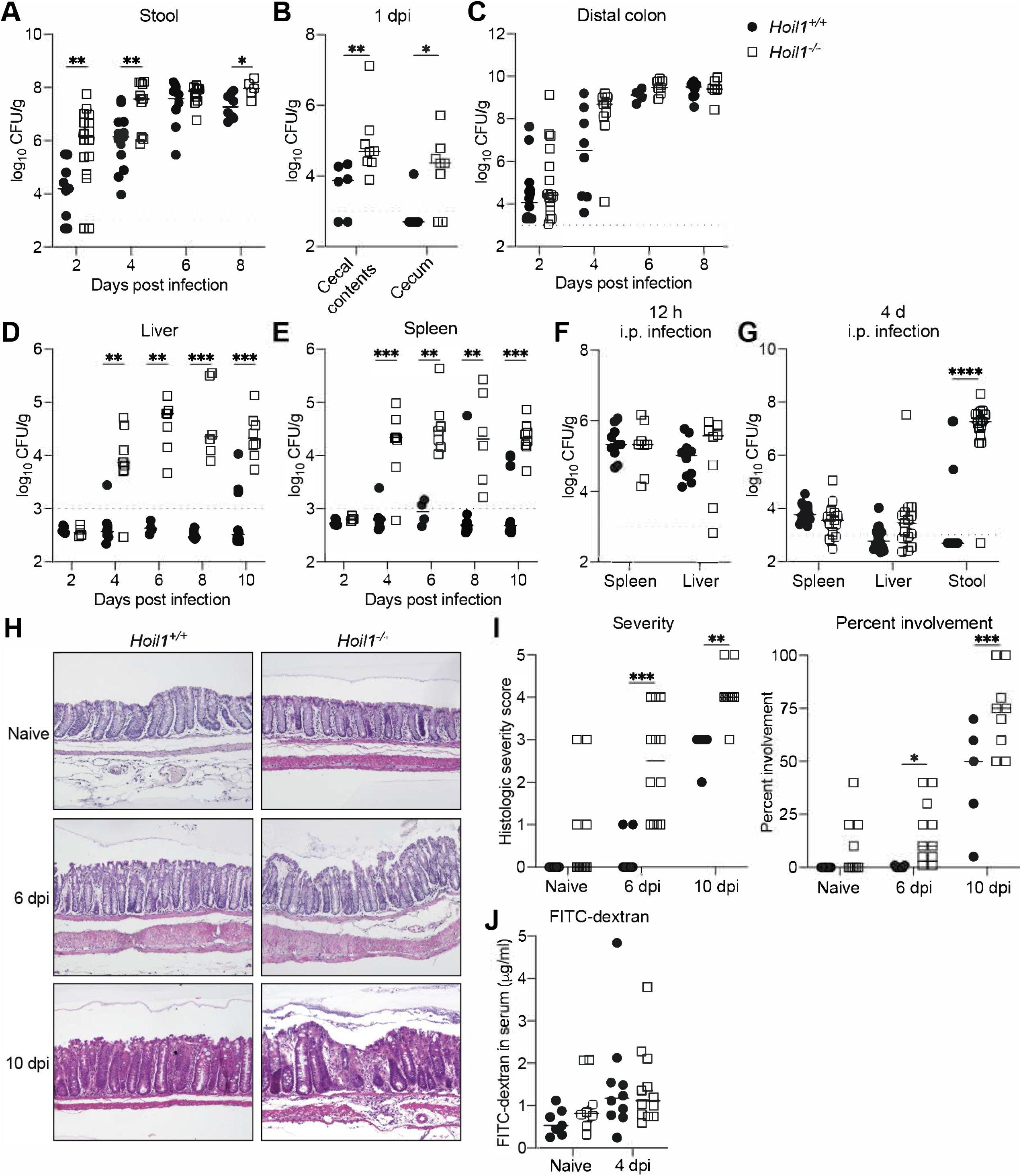
HOIL1 is essential to prevent bacterial dissemination to systemic sites and colonic ulceration following *C. rodentium* infection. **A-E.** *C. rodentium* CFU in stool (A), cecal contents and cecum (B), distal colon (C), liver (D) and spleen (E) of *Hoil1*^+/+^ (circles) and *Hoil1*^-/-^ (squares) mice on indicated days following intragastric (i.g.) inoculation with 2x10^9^ CFU *C. rodentium*. **F**. *C. rodentium* CFU in spleen and liver 12 h post-infection of *Hoil1^+/+^* and *Hoil1^-/-^*mice following intraperitoneal (i.p) inoculation with 10^7^ CFU. **G**. *C. rodentium* CFU in spleen, liver, and stool of *Hoil1^+/+^* and *Hoil1^-/-^* mice 4 d following i.p. inoculation with 10^7^ CFU. **H**. Representative H&E-stained sections of distal colon from naïve *Hoil1^+/+^* (left panels) and *Hoil1^-/-^* (right panels) mice (top), or mice inoculated i.g. with *C. rodentium* for 6 d (middle) or 10 d (bottom). **I**. Histological severity score (left) and percent of colon length involvement (right) of *Hoil1^+/+^* (circles) and *Hoil1^-/-^*(squares) mice following i.g. infection with *C. rodentium* at 0, 6, and 10 d post-infection. Histological scoring criteria: 0, no neutrophilic inflammation; 1, neutrophilic inflammation in surface epithelium, lamina propria, or cryptitis; 2, crypt abscess; 3, neutrophilic inflammation present at muscularis mucosae or beyond; 4, ulcer or transmural inflammation; 5, epithelium denudement. **J**. Concentration of FITC-dextran in serum 4 h after i.g. administration in *Hoil1^+/+^* and *Hoil1^-/-^* naïve mice, and at 4 d post-inoculation with 2x10^9^ CFU *C. rodentium* i.g. Each symbol represents an individual mouse. Dotted line indicates limit of detection (LOD). Data were combined from at least 3 independent experiments. Statistical analyses were performed using Mann-Whitney test. \**p*≤0.05, \*\**p*≤0.01, \*\*\**p*≤0.001.

We considered that the detection of *C. rodentium* CFU in the spleen and liver of *Hoil1*^-/-^ mice could be due to increased dissemination of bacteria from the intestine, or due to failure to kill bacterial that have reached these sites. To determine whether HOIL1 is required to control *C. rodentium* replication at systemic sites, we by-passed the intestinal epithelial barrier by infecting mice intraperitoneally with 10^7^ CFU *C. rodentium.* Burdens in the spleen and liver were comparable between *Hoil1^+/+^* and *Hoil1*^-/-^ mice at 12 hpi and at 4 dpi (Fig. 1F-G), suggesting that HOIL1 is not required to control *C. rodentium* replication outside of the intestine. However, by 4 dpi, *Hoil1*^-/-^ mice infected intraperitoneally were shedding large amounts of *C. rodentium* in the stool indicating that infection of the GI tract had occurred in these mice (Fig. 1G), and that later time points would be uninformative. Together, these data indicate that HOIL1 plays an important role in restricting intestinal replication and attachment of *C. rodentium*, as well as in preventing its dissemination to systemic organs.

Since the previous experiments suggested that HOIL1 regulates intestinal barrier function, we next examined the severity of intestinal pathology and colitis over the course of infection. Thirty percent of the uninfected *Hoil1^-/-^* mice exhibited neutrophils in the epithelium, lamina propria, or muscularis mucosae (Fig. 1H and 1I). By 6 dpi, all *Hoil1^-/-^* mice exhibited inflammation, and 25% of these mice had developed ulcers, whereas only superficial neutrophilic inflammation was observed in the colon of 25% of *Hoil1^+/+^* mice. By 10 dpi, *Hoil1^+/+^* mice developed characteristic acute colitis, consisting of increased crypt depth, depletion of goblet cells, infiltration of immune cells and edema. In contrast, HOIL1-mutant mice developed severe colitis, including ulceration, transmural inflammation and, in some cases, epithelial denudement. Additionally, a larger proportion of the colon was inflamed in *Hoil1^-/-^* mice at both time points (Fig. 1I). These data indicate that HOIL1 expression is essential to protect against excessive inflammation and ulceration of the colon during *C. rodentium* infection.

We next asked whether HOIL1-deficiency resulted in an increase in intestinal permeability. Since *C. rodentium* was detected in systemic organs of *Hoil1^-/-^* mice by 4 dpi, we expected that any changes in intestinal permeability responsible for bacterial translocation would be detectable by this time. Naïve mice, or mice infected for 4 days were administered FITC-dextran (4000 average molecular weight) by oral gavage, and the concentration of FITC in the serum measured 4 hr later (Fig. 1J). However, we did not observe a significant difference in the amount of FITC-dextran present in the serum of *Hoil1^+/+^* and *Hoil1^-/-^* mice before or during *C. rodentium* infection, suggesting that HOIL1 may not be required to regulate intestinal permeability in the early phases of *C. rodentium* infection.

Together, these data indicate that HOIL1 is essential to limit colonic inflammation and prevent dissemination of *C. rodentium* to systemic organs following infection. However, HOIL1 does not appear to regulate passive intestinal permeability at 4 dpi, at which time *C. rodentium* CFU are detectable in systemic organs, suggesting that HOIL1 prevents dissemination through an active mechanism.

### HOIL1 modulates the induction of a subset of cytokines in the colon during *C. rodentium* infection

We considered that HOIL1 deficiency might lead to defects or delays in induction of inflammatory cytokines and anti-microbial mediators during *C. rodentium* infection that allow for the bacteria to replicate and colonize the cecum and colon more rapidly (Fig. 1A-C). We have shown previously that HOIL1-mutant mice mount a defective type 1 inflammatory response, including impaired induction of IL-12, IFNγ and iNOS, following *L. monocytogenes* infection (24). Since type 3 inflammatory cytokines are an essential component of the mucosal innate immune response to *C. rodentium* (17, 49), and IL-12 and IL-23 share subunit IL-12p40, we considered that HOIL1 may be required for induction of IL-23, and subsequently IL-22 and downstream responses. We therefore measured cytokine mRNA and protein levels in the distal colon over the course of infection. However, *Il12b* mRNA levels were not significantly different during 8 days of infection (Fig. 2A), and we were not able to detect IL-12p40 protein in the colon of naïve *Hoil1^+/+^*or *Hoil1^-/-^* mice or at 6 dpi (not shown). Similarly, *Il23a* mRNA levels changed minimally in both *Hoil1^+/+^* and *Hoil1^-/-^* mice over the course of infection. IL-23 protein was slightly elevated in naïve *Hoil1^-/-^* distal colon, and slightly, but not significantly lower at 6 dpi compared to *Hoil1^+/+^* colon (Fig. 2A). IL-1β protein expression was similar in *Hoil1*^+/+^ and *Hoil1^-/-^* tissue from naïve mice and at 6 dpi (Fig. 2B). We next examined the induction of IL-22, which is produced by ILC3 and Th22 cells in response to IL-23 and IL-1β (17). At 6 dpi, *Il22* mRNA was approximately three-fold lower, and IL-22 protein was two-fold lower in *Hoil1^-/-^* tissue relative to *Hoil1^+/+^* tissue (Fig. 2C). Since *C. rodentium* burdens in the cecum were elevated by 1 dpi, we also examined cytokine expression in the cecum early during infection. However, both *Il23a* (Fig. 2D) and *Il22* (Fig. 2E) were poorly induced in the cecum at 1 and 2 dpi, and mRNA levels were similar for *Hoil1^+/+^* and *Hoil1^-/-^* mice.

**Figure 2:**
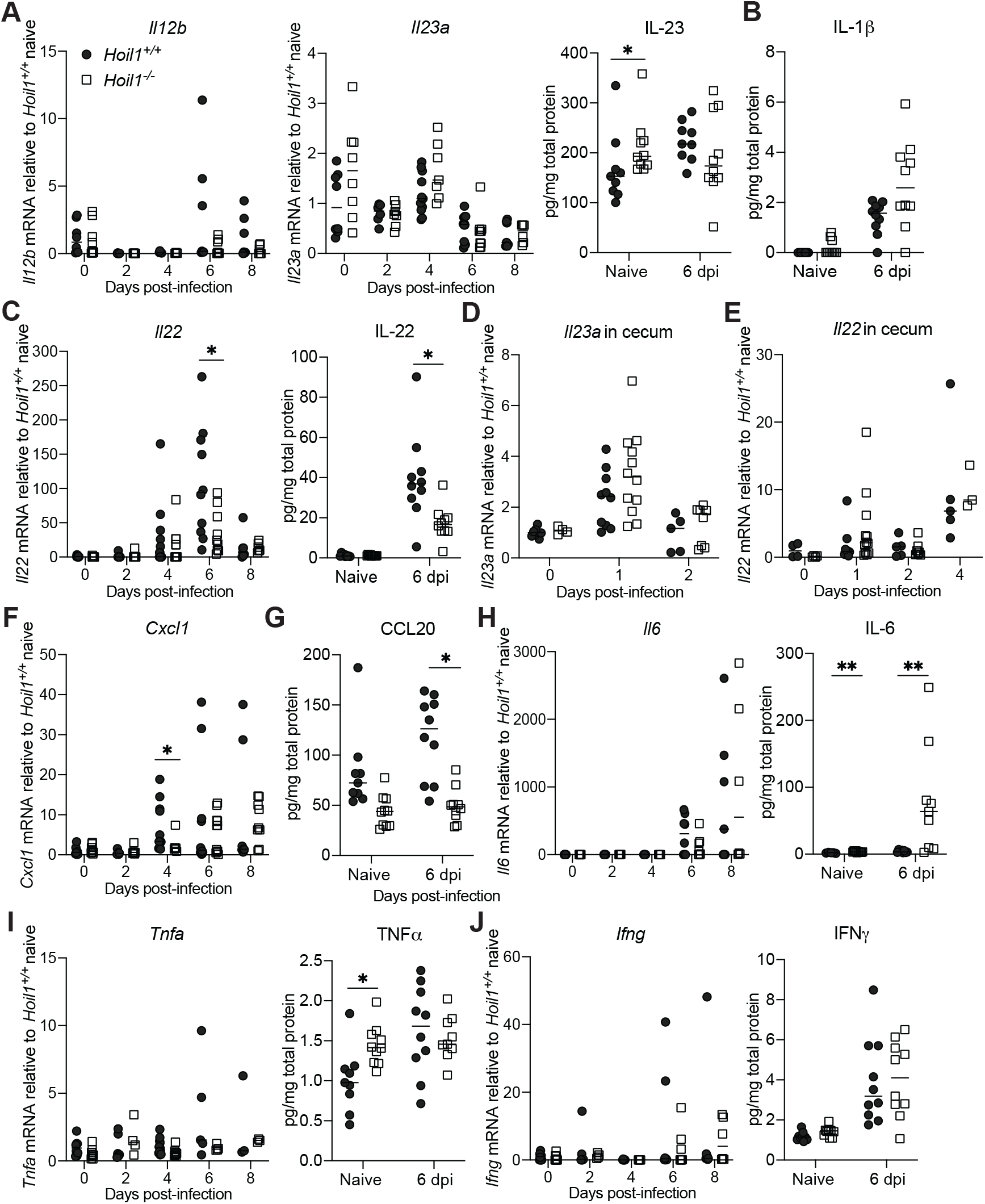
HOIL1 modulates the induction of a subset of cytokines in the colon during *C. rodentium* infection. Relative cytokine mRNA levels measured by RT-qPCR in the distal colon over 8 days in *Hoil1^+/+^* (circles) and *Hoil1^-/-^* (squares) mice following i.g. infection with 2x10^9^ CFU *C. rodentium,* or protein concentrations in homogenized colon from *Hoil1^+/+^* and *Hoil1^-/-^* mice, naïve and at 6 dpi. **A**. *Il12b* mRNA (left), *Il23a* mRNA (center), IL-23 protein (right). **B**. IL-1β protein. **C**. *Il22* mRNA (left) and IL-22 protein (right). **D**. *Il23a* mRNA in cecum at 0, 1, and 2 dpi. **E**. *Il22* mRNA in cecum at 0, 1, and 2 dpi. **F**. *Cxcl1* mRNA. **G.** CCL20 protein. **H**. *Il6* mRNA (left) and IL-6 protein (right). **I**. *Tnfa* mRNA (left) and TNF⍺ protein (right) **J**. *Ifng* mRNA (left) and IFNγ protein (right). Each symbol represents an individual mouse. Data were combined from at least three independent experiments. Statistical analyses performed by Mann-Whitney test. \**p*≤0.05, \*\**p*≤0.01.

Chemokines are important for the recruitment of innate immune effector cells such as neutrophils and inflammatory monocytes to the site of infection. *Cxcl1* mRNA was lower in *Hoil1^-/-^* mice at 4 dpi (Fig. 2F), and CCL20 protein was slightly lower at baseline and significantly reduced during infection (Fig. 2G) compared to *Hoil1^+/+^* mice, suggesting that recruitment of effector cells could be impaired.

Since histological examination had indicated increased ulceration in *Hoil1^-/-^* colon during infection (Fig. 1H-I), we considered that an excessive type 1 inflammatory response might be responsible. Although *Il6* mRNA was induced similarly in *Hoil1^+/+^* and *Hoil1^-/-^* colon tissue, IL-6 protein was significantly higher in *Hoil1^-/-^*tissue at baseline and at 6 dpi (Fig. 2H). *Tnfa* mRNA was poorly induced and differences between genotypes were not significant during infection, although TNF-α protein was elevated in naïve *Hoil1^-/-^* tissue (Fig 2I). Although *Ifng* mRNA levels appeared to be slightly higher in *Hoil1^-/-^* tissue at 6 and 8 dpi, protein levels were induced similarly at 6 dpi (Fig 2J).

Overall, we observed a reduction in IL-22 and CCL20 production during infection, as well as a slight increase in IL-6 at later stages of infection that could be either causative or a consequence of the pathological changes to the tissue. These data indicate that HOIL1 modulates the balance of inflammatory cytokines in the colon during infection.

### HOIL1 plays an important role in the innate immune response to *C. rodentium*

We next sought to determine which cells require HOIL1 expression to prevent weight loss, systemic dissemination, and lethality following *C. rodentium* infection. Both the innate and adaptive branches of the immune response are required to control, clear, and survive *C. rodentium* infection (17, 23). To test whether HOIL1 plays a predominant role in innate or adaptive immune responses, we challenged *Rag1^-/-^Hoil1^+/+^*and *Rag1^-/-^Hoil1^-/-^* mice lacking mature B and T cells with *C. rodentium*. As reported by others (23), *Rag1^-/-^Hoil1^+/+^* mice began to lose weight around 12 dpi and succumbed to the infection between 18 and 22 dpi (Fig. 3A-B, top). However, *Rag1^-/-^Hoil1^-/-^* mice lost weight and succumbed to the infection approximately 7 days earlier, and in a similar time frame to *Hoil1^-/-^* mice (Fig. 3A-B, bottom; (24)). Furthermore, *Rag1^-/-^Hoil1^-/-^*mice shed significantly more *C. rodentium* in their stool than *Rag1^-/-^Hoil1^+/+^* mice as early as 2 dpi (Fig. 3C), and exhibited higher titers in the colon and cecum at 6 dpi (Fig. 3D), similar to *Hoil1^-/-^* mice (Fig. 1). By 6 dpi, *C. rodentium* was detected at systemic sites including the liver, spleen, and mesenteric lymph nodes (MLN) in *Rag1^-/-^Hoil1^-/-^* mice (Fig. 3D), while bacteria were rarely found outside the colon and cecum in *Rag1^-/-^Hoil1^+/+^* mice at this time point. Although these data do not rule out a role for HOIL1 in the adaptive immune response to *C. rodentium*, they demonstrate that HOIL1 plays an essential role in the early innate immune response to *C. rodentium* in the absence of adaptive immunity.

**Figure 3:**
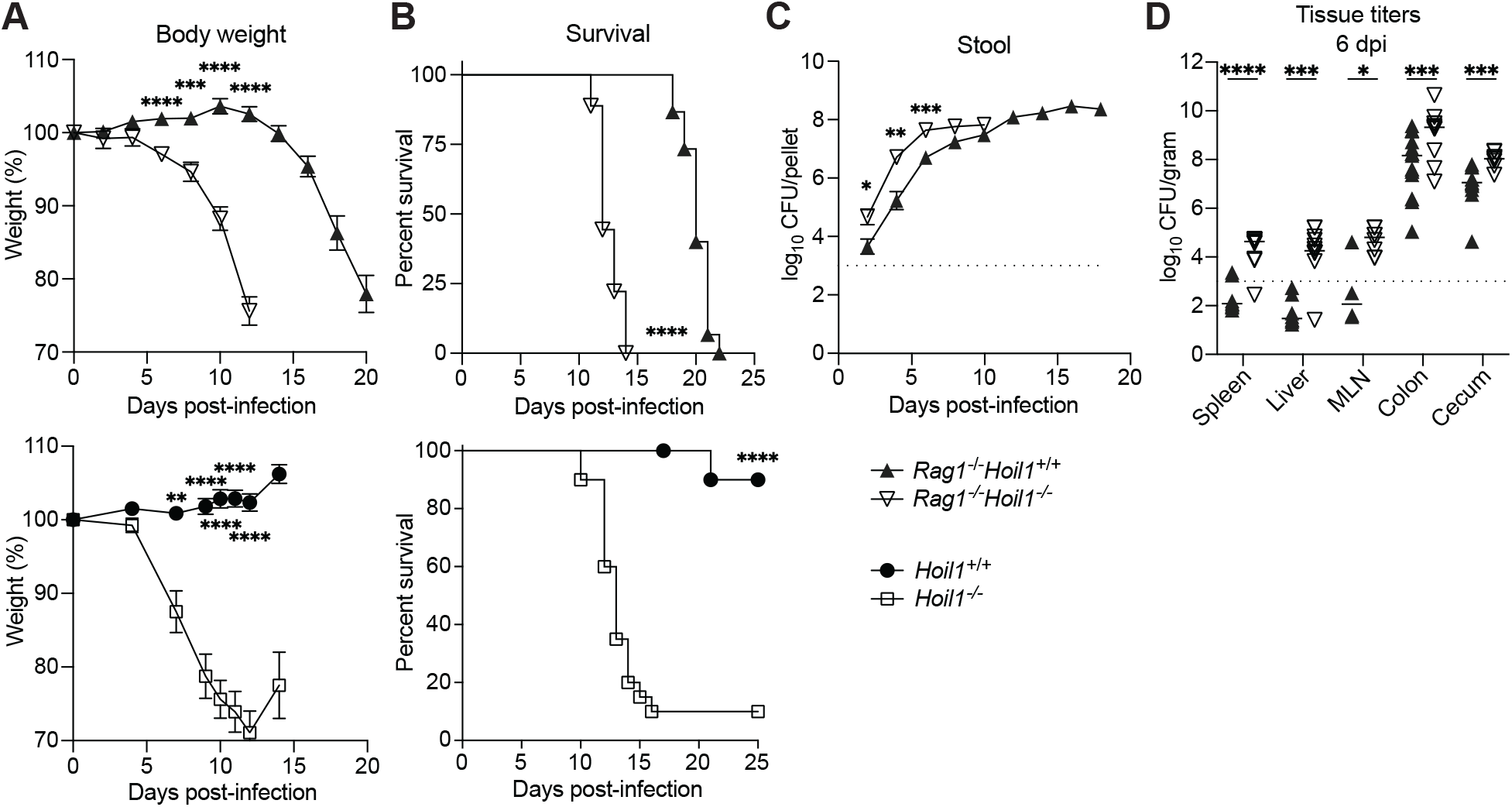
HOIL1 plays an essential function in innate immunity during *C. rodentium* infection. **A**. Percent initial body weight of *Rag^-/-^Hoil1^+/+^*(filled triangles; n=15) and *Rag^-/-^Hoil1^-/-^* (inverted triangles; n=9) (top panel), and *Hoil1^+/+^* (filled circles; n=10) and *Hoil1^-/-^* (squares; n=10) (bottom panel) mice at the indicated times following i.g. inoculation with 2x10^9^ CFU *C. rodentium*. **B**. Survival of *Rag^-/-^Hoil1^+/+^*(n=15) and *Rag^-/-^Hoil1^-/-^* (n=9) (top panel), and *Hoil1^+/+^* (n=20) and *Hoil1^-/-^* (n=20) (bottom panel) mice following i.g. inoculation with 2x10^9^ CFU *C. rodentium*. **C**. *C. rodentium* CFU in stool from *Rag^-/-^Hoil1^+/+^* (n=15) and *Rag^-/-^Hoil1^-/-^* (n=9) mice at the indicated times following i.g. inoculation with 2x10^9^ CFU *C. rodentium*. Data shown indicate the mean ± standard error. **D**. *C. rodentium* CFU in spleen, liver, mesenteric lymph node (MLN), distal colon and cecum of *Rag^-/-^Hoil1^+/+^* and *Rag^-/-^Hoil1^-/-^*6 dpi with 2x10^9^ CFU *C. rodentium* i.g. Each symbol represents an individual mouse and horizontal bars indicate the median. Data in bottom panels of (A) and (B) reprinted from MacDuff *et. al*., eLife 2015 (24). Dotted lines indicate the LOD. Data were combined from at least three independent experiments. Statistical analyses performed using unpaired *t*-test with Welch correction (A), Mann-Whitney test (C, D) or log-rank Mantel-Cox test (B). \**p*≤0.05, \*\**p*≤0.01, \*\*\**p*≤0.001, \*\*\*\**p*≤0.0001.

### Expression of HOIL1 in radiation-resistant cells is important during *C. rodentium* infection

Both hematopoietic and non-hematopoietic cells, such as intestinal epithelial cells (IEC), are critical for the immune response to *C. rodentium* in the intestine (23). To determine if HOIL1 is required in hematopoietic or non-hematopoietic cells, we generated reciprocal bone marrow chimeric mice and allowed 16 weeks for bone marrow reconstitution and recovery from irradiation. Chimerism was confirmed by measuring the percent wild-type (WT) and knock-out (KO) gDNA in the blood (Fig. 4A). These mice were infected with *C. rodentium* and monitored for weight loss over 10 days (Fig. 4B). WT mice reconstituted with WT bone marrow (WT->WT) lost approximately eight percent of their body weight over this time, indicating that 16 weeks was not sufficient for full recovery from irradiation. KO mice that received KO bone marrow (KO->KO) lost the greatest percentage of body weight (approximately 17%), as expected. Surprisingly, WT mice that received KO bone marrow (KO->WT) progressed similarly to WT mice that received WT bone marrow, indicating that loss of HOIL1 in bone marrow-derived cells does not significantly increase susceptibility to *C. rodentium-*induced morbidity by 10 dpi. KO mice that received WT bone marrow (WT->KO) displayed an intermediate phenotype, with delayed weight loss compared to KO->KO mice, but ultimately lost a similar percentage of body weight by 10 dpi. Thus, HOIL1 plays an important role in radiation-resistant cells during *C. rodentium* infection, and expression of HOIL1 in hematopoietic cells is only partially protective. These findings were further supported by increased *C. rodentium* CFU in the liver, and more severe pathology observed in colon from WT->KO mice at 10 dpi (Fig. 4C, D). However, the irradiated mice developed more severe pathology than non-irradiated mice (Fig. 4D and Fig. 2B), indicating that the recovery from irradiation may be a confounding factor. Together, these data support a major role for HOIL1 in radiation-resistant to limit disease severity during *C. rodentium* infection, and a more minor role for HOIL1 in hematopoietic cells.

**Figure 4:**
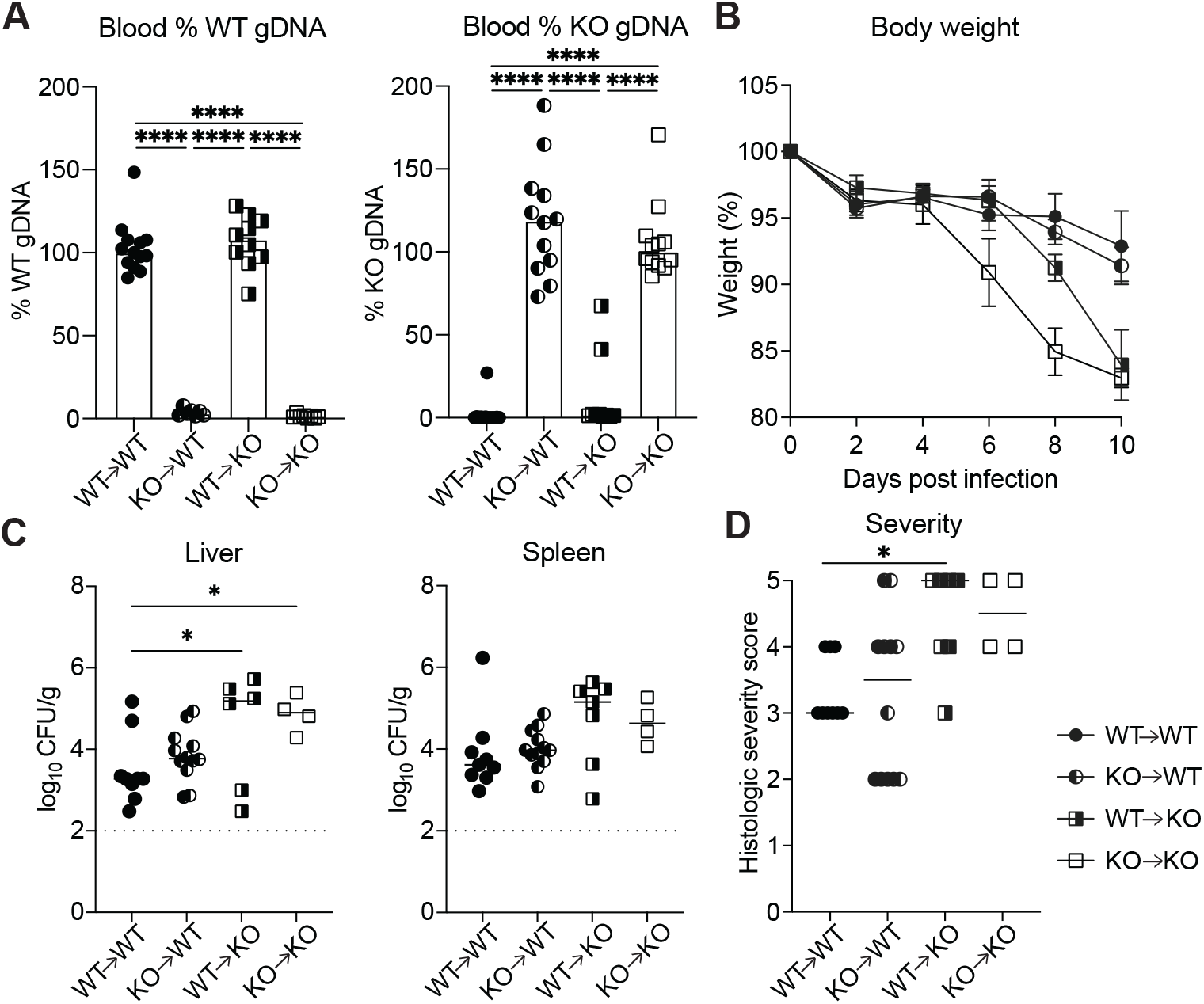
Expression of HOIL1 is important in radiation-resistant cells during *C. rodentium* infection. **A.** Percent of WT (*Rbck1* intron 7, left panel) or KO (Neomycin-resistance gene, right panel) gDNA in blood from bone marrow chimeric mice relative to WT->WT and KO->KO controls. Each symbol represents and individual mouse and bars indicate the median. **B**. Percent of initial weight of *Hoil1* WT->WT (n=9), *Hoil1* KO->WT (n=12), *Hoil1* WT->KO (n=8) and *Hoil1* KO->KO (n=8) bone marrow chimeric mice over 10 days following i.g. inoculation with 2x10^9^ CFU *C. rodentium.* Data represent the mean +/- standard error. Significant differences at 8 dpi: WT->WT vs KO->KO, KO->WT vs KO->KO, and WT->KO vs KO->KO. Significant differences at 10 dpi: WT->WT vs KO->KO, and KO->WT vs KO->KO. **C.** *C. rodentium* CFU in liver and spleen at 10 dpi. **D.** Histologic severity score of colon at 10 dpi. For A, C, D, each symbol represents an individual mouse. Data were combined from at least 3 independent experiments. Statistical analyses performed by Brown-Forsythe and Welch one-way ANOVA with Dunnett’s T3 multiple comparisons test (A and D), one-way ANOVA relative to WT->WT (C), or by mixed effects analysis with Tukey’s multiple comparisons test (B). \**p*≤0.05, \*\**p*≤0.01, \*\*\**p*≤0.001, \*\*\*\**p*≤0.0001.

### HOIL1 is important in CD11c- and lysozyme-2-expressing myeloid cells to prevent systemic dissemination of *C. rodentium*

To further identify which cell types require HOIL1 expression, we generated conditional knockout mice. Knock-out first mice with conditional potential were acquired from the European Mutant Mouse Archive (EMMA), and used to generate mice with exons 5 and 6 of the *Rbck1* gene, which encodes the HOIL1 protein, flanked by loxP sites. Deletion of these exons by crossing to mice expressing cre recombinase driven by the CMV promoter resulted in viable *Rbck1^-/-^* mice (herein referred to as such to distinguish them from the original *Hoil1^-/-^* mouse line), indicating that these mice are not fully LUBAC deficient similar to the *Hoil1^-/-^* mice. Indeed, HOIP and SHARPIN protein levels were reduced, but not completely absent in the spleen from *Rbck1*^-/-^ mice (Fig. S1A). *Rbck1* mRNA levels upstream of the deleted region were similar to wild-type levels (Fig. S1B), indicating that the N-terminal region containing the UBL domain important for complex formation may be expressed (43).

To confirm that *Rbck1^-/-^*mice displayed a similar phenotype to *Hoil1^-/-^* mice, we infected *Rbck1^f/f^* and *Rbck1^-/-^* mice with *C. rodentium* and monitored them for weight loss and survival. *C. rodentium* infection resulted in weight loss beginning around 6 dpi, and lethality in 57% of *Rbck1^-/-^* mice beginning around 10 dpi (Fig. S1C-D), similar to *Hoil1^-/-^* mice (Fig. 3) (24). Weight loss in *Rbck1^-/-^* mice was slightly delayed compared to *Hoil1^-/-^* mice, however, and surviving mice recovered their body weight by 20 dpi. *Rbck1^-/-^* mice shed more *C. rodentium* in their stool early in infection (Fig. S1E), and their colons were colonized more rapidly by *C. rodentium* compared to *Rbck1^f/f^*mice (Fig. S1F). Furthermore, *C. rodentium* spread to systemic organs including the liver and spleen in *Rbck1^-/-^* mice (Fig. S1G-H), although with slightly delayed kinetics compared to *Hoil1^-/-^* mice (Fig. 1D, E). Overall, these data indicate that *Rbck1^-/-^* mice have a similar phenotype to *Hoil1^-/-^* mice and are an appropriate model to examine the role of HOIL1 in specific cell types.

Since the susceptibility of the bone marrow chimeric mice to infection indicated that a radiation-resistant cell type was important, we hypothesized that HOIL1 expression may be important in intestinal epithelial cells (IECs), which are the primary site of *C. rodentium* infection. However, no differences were observed in the weight loss or bacterial shedding in the stool of *Rbck1^f/-^* and *Rbck1^f/-^* ^Vil1cre^ littermates over the course of infection (Fig. 5A-B), and the colon was colonized to similar levels at 6 and 8 dpi (Fig. 5C). *C. rodentium* CFU were rarely detected in the liver, spleen, or MLN of *Rbck1^f/-^* or *Rbck1^f/-^* ^Vil1-cre^ mice at 6 or 8 dpi (Table 1). We confirmed efficient deletion of *Rbck1* in IEC in *Rbck1^f/-^*^Vil1cre^ mice by qRT-PCR (Fig. S1I). These data indicate that HOIL1 is not required in IECs to control *C. rodentium* infection.

**Figure 5:**
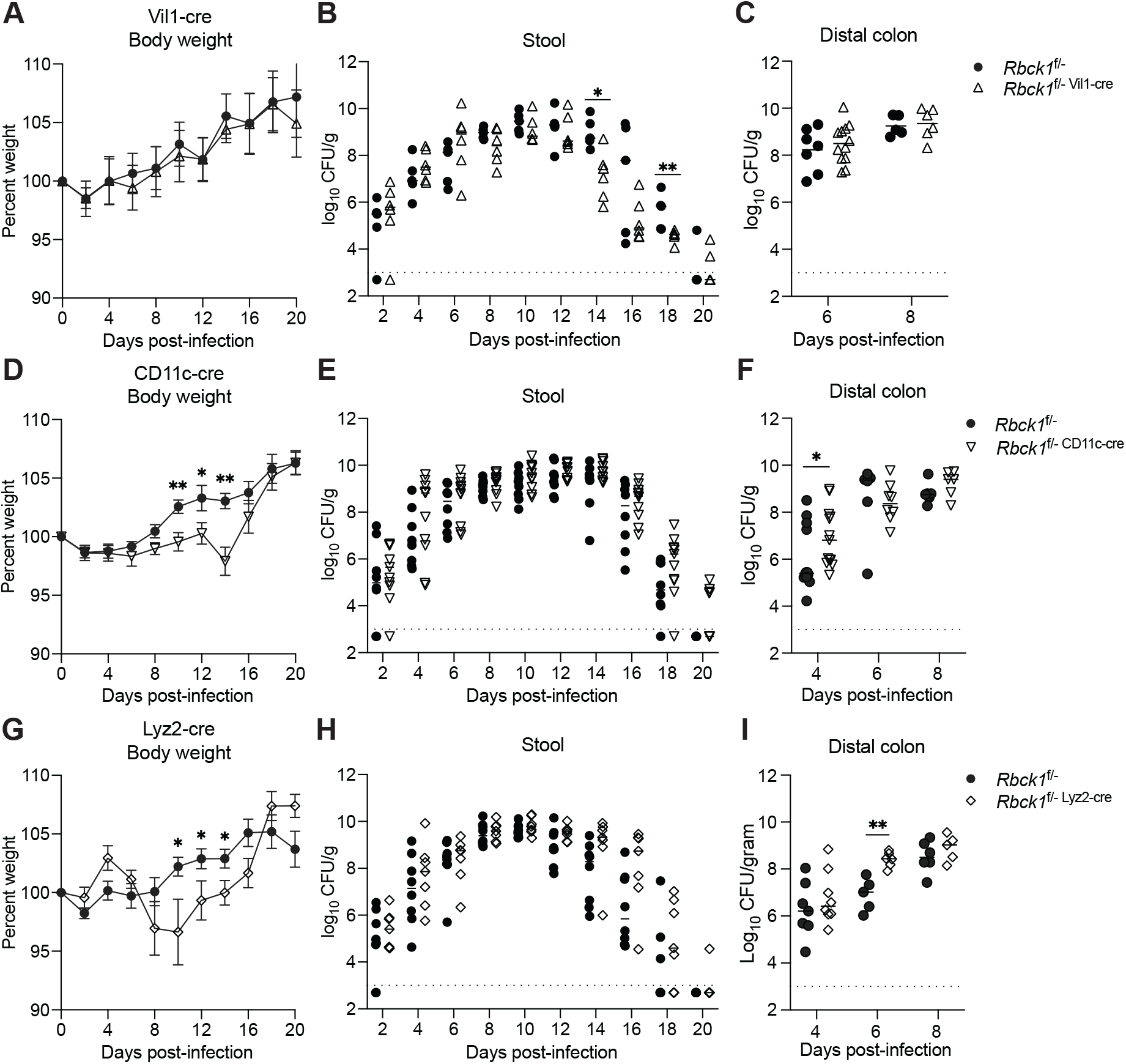
HOIL1 functions in CD11c- and lysozyme 2-expressing myeloid cells to prevent weight loss and systemic dissemination following *C. rodentium* infection. **A-B.** Percent body weight (A) and *C. rodentium* CFU in stool (B) from *Rbck1*^f/-^ (circles, n=5) and *Rbck1*^f/-Vil1-cre^ (triangles, n=6) mice at the indicated times following i.g. inoculation. **C**. *C. rodentium* CFU in distal colon from *Rbck1*^f/-^ and *Rbck1*^f/-Vil1-cre^ mice at 6 and 8 dpi. D-E. Percent body weight (D) and *C. rodentium* CFU in stool (E) of *Rbck1*^f/-^ (circles, n=8) and *Rbck1*^f/-CD11c-cre^ (inverted triangles, n=11) mice at the indicated times following i.g. inoculation. F. *C. rodentium* CFU in distal colon from *Rbck1*^f/-^ and *Rbck1*^f/-CD11c-cre^ mice at 4, 6, and 8 dpi. **G-H**. Percent body weight (G) and *C. rodentium* CFU in stool (H) of *Rbck1*^f/-^ (circles, n=11) and *Rbck1*^f/-Lyz2-cre^ (diamonds, n=10) mice at the indicated times following i.g. inoculation. I. *C. rodentium* CFU in distal colon from *Rbck1*^f/-^ and *Rbck1*^f/-Lyz2-cre^ mice at 4, 6, and 8 dpi. Each symbol represents an individual mouse. Dotted lines indicate the LOD. Data were combined from at least three independent experiments. Statistical analyses performed by unpaired t-test with Welch correction (A, D, G) or Mann-Whitney test (B, C, E, F, H, I). \**p*≤0.05, \*\**p*≤0.01.

**Table 1:** Proportion of *Rbck1*^f/-^, *Rbck1*^f/-Vil1-cre^, *Rbck1*^f/-CD11c-cre^, and *Rbck1*^f/-Lyz2-cre^ mice with systemic dissemination following *C. rodentium* infection. *C. rodentium* CFU in spleen, liver, and MLN from *Rbck1*^f/-Vil1-cre^, *Rbck1*^f/-CD11c-cre^, *Rbck1*^f/-Lyz2-cre^ and respective *Rbck1*^f/-^ littermate control mice at the indicated time points. Results are presented as the number of mice with detectable *C. rodentium* CFU / total number of mice evaluated (left columns), and the percentage of mice with detectable CFU (right columns). MLN; mesenteric lymph nodes.

|  | Liver |  |  |  | Spleen |  |  |  | MLN |  |  |  |
| --- | --- | --- | --- | --- | --- | --- | --- | --- | --- | --- | --- | --- |
|  | proportion | percent | proportion | percent | proportion | percent | proportion | percent | proportion | percent | proportion | percent |
|  | *Rbck1*<sup>f/-</sup> |  | f/- +CD11c cre |  | *Rbck1*<sup>f/-</sup> |  | f/- +CD11c cre |  | *Rbck1*<sup>f/-</sup> |  | f/- +CD11c cre |  |
| D4 | 0/12 | 0 | 3/13 | 23.1 | 0/12 | 0.0 | 3/13 | 23.1 | 0/12 | 0.0 | 4/13 | 30.8 |
| D6 | 0/6 | 0 | 3/8 | 37.5 | 0/6 | 0.0 | 3/8 | 37.5 | 0/6 | 0.0 | 3/8 | 37.5 |
| D8 | 0/11 | 0 | 6/15 | 40.0 | 1/11 | 9.1 | 7/15 | 46.7 | 1/11 | 9.1 | 5/15 | 33.3 |
|  | *Rbck1*<sup>f/-</sup> |  | f/- +LysM cre |  | *Rbck1*<sup>f/-</sup> |  | f/- +LysM cre |  | *Rbck1*<sup>f/-</sup> |  | f/- +LysM cre |  |
| D4 | 0/7 | 0 | 1/8 | 12.5 | 0/7 | 0.0 | 1/8 | 12.5 | 0/7 | 0.0 | 1/8 | 12.5 |
| D6 | 0/5 | 0 | 2/6 | 33.3 | 0/5 | 0.0 | 2/6 | 33.3 | 0/5 | 0.0 | 1/6 | 16.7 |
| D8 | 0/10 | 0 | 5/8 | 62.5 | 0/10 | 0.0 | 5/8 | 62.5 | 0/10 | 0.0 | 2/8 | 25.0 |
|  | *Rbck1*<sup>f/-</sup> |  | f/- +Vil1 cre |  | *Rbck1*<sup>f/-</sup> |  | f/- +Vil1 cre |  | *Rbck1*<sup>f/-</sup> |  | f/- +Vil1 cre |  |
| D4 | ND | ND | ND | ND | ND | ND | ND | ND | ND | ND | ND | ND |
| D6 | 0/7 | 0 | 1/12 | 8.3 | 0/7 | 0.0 | 1/12 | 8.3 | 0/7 | 0.0 | 1/12 | 8.3 |
| D8 | 0/7 | 0 | 0/8 | 0.0 | 0/7 | 0.0 | 1/8 | 12.5 | 1/7 | 14.3 | 0/8 | 0.0 |

Proportion and percent mice of the indicated genotypes with *C. rodentium* detected in the liver, spleen and MLN at 4, 6 and 8 dpi. *Rbck1*<sup>f/-</sup> littermate controls were used for each of the cre-expressing strains.

Dendritic cells (DCs) and macrophages are a critical early responder during *C. rodentium* infection, producing cytokines such as IL-23 that is important to initiate the immune response and limit pathogenesis during infection (14, 17, 50–52). Since we observed a defect in IL-22 induction (Fig. 2C), we wondered if HOIL1 is required in CD11c^+^ mononuclear phagocytes to prevent severe pathology following *C. rodentium* infection. Compared to *Rbck1^f/-^*littermates, *Rbck1^f/-^* ^CD11c-cre^ mice lost more body weight starting around 6-8 dpi and peaking at 14 dpi, but recovered by 18 dpi (Fig. 5D). No difference was observed in bacterial shedding in the stool (Fig. 5E), but *C. rodentium* colonized the colon more rapidly in *Rbck1^f/-^*^CD11c-cre^ mice compared to *Rbck1^f/-^* mice, leading to an increase in bacterial titers at 4 dpi (Fig. 5F). Furthermore, *C. rodentium* CFU were detected in systemic organs of 23% of *Rbck1^f/-^* ^CD11c-cre^ mice by 4 dpi and approximately 40% by 8 dpi (Table 1). These data indicate that HOIL1 plays an important role in CD11c^+^ mononuclear phagocytes to prevent systemic spread of *C. rodentium* and weight loss during infection.

We also examined the role of HOIL1 in lysozyme 2-expressing cells, which include macrophages and neutrophils that may be important for bacterial phagocytosis and clearance. *Rbck1^f/-^*^Lyz2-cre^ mice lost weight relative to their *Rbck1^f/-^*littermates starting around 8 dpi and peaking at 10 dpi, but recovered by 18 dpi (Fig. 5G). *Rbck1^f/-^* ^Lyz2-cre^ mice and *Rbck1^f/-^* mice shed similar amounts of bacteria early in infection (Fig. 5H) and *C. rodentium* burdens were similar in intestinal tissue (Fig. 5I). We detected *C. rodentium* CFU in systemic organs of approximately 62.5% of *Rbck1^f/-^* ^Lyz2-cre^ mice at 8 dpi (Table 1). These data indicate that HOIL1 plays a role in lysozyme 2-expressing cells to control *C. rodentium* dissemination and limit morbidity.

Overall, these data indicate that HOIL1 functions in CD11c- and lysozyme-expressing myeloid cells to prevent morbidity and systemic dissemination of bacteria following *C. rodentium* infection.

### HOIL1 is important for maintenance of ILC3 numbers in the colon

To investigate whether HOIL1 regulates the maintenance or recruitment of CD11c- and Lyz2-expressing cells to the colon, we characterized immune cell populations in the colonic lamina propria in naïve mice and at 6 dpi with *C. rodentium*. However, proportions of CD11b^+^ Ly6G^+^ neutrophils, CD11c^+^ CD11b^+^ CX3CR1^+^ resident macrophages, and CD11b^+^ Ly6C^+^ monocytes were similar in the lamina propria from *Hoil1*^+/+^ and *Hoil1*^-/-^ mice before infection and at 6 dpi (Fig. 6A-C, Fig. S2A-C). CD11c^+^ CD103^+^ CD11b^+^ DCs that have been shown to produce IL-23 were also unaffected by HOIL1-deficiency, but there were significantly fewer CD11c^+^ CD103^+^ CD11b^-^ DCs that have been shown to produce IL-12, and total CD11c^+^ CD103^+^ dendritic cells in the lamina propria from *Hoil1*^-/-^ mice at 6 dpi (Fig. 6D, Fig. S2A, D) (14, 51–54). These data indicate that HOIL1 is not required to control populations of neutrophils, resident macrophages, or monocytes, but may be important for the maintenance or recruitment of a subset of DCs during *C. rodentium* infection.

**Figure 6:**
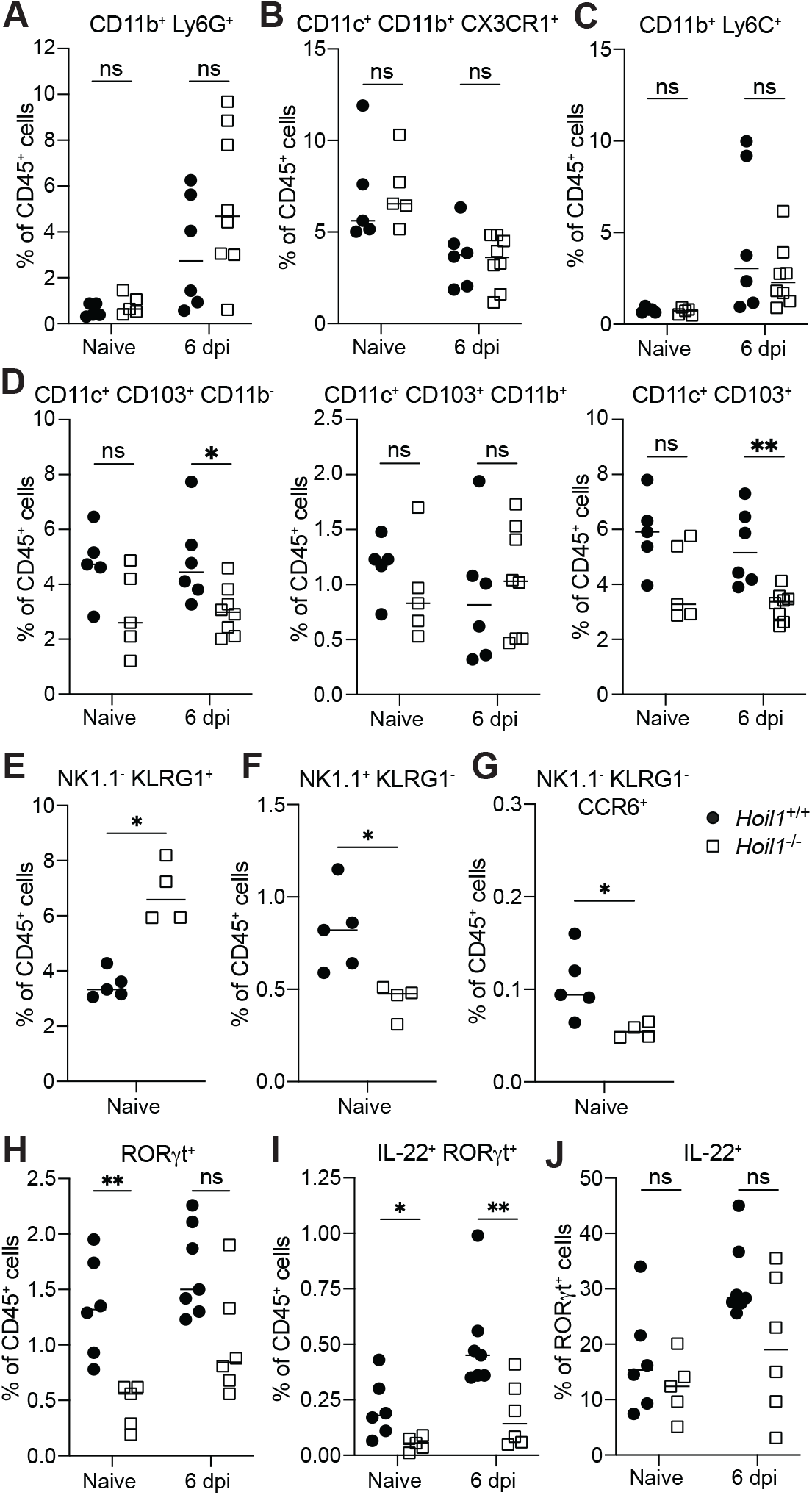
HOIL1 is important for maintenance of ILC3 numbers in the colon. Quantification of innate immune cell populations in the colonic lamina propria from naïve *Hoil1^+/+^*(left) and *Hoil1^-/-^* (right) mice and at 6 dpi with *C. rodentium.* Pre-gated on live, CD45^+^ singlets, and presented as the percent of CD45^+^ cells (except J). **A**. Ly6G^+^ CD11b^+^ (neutrophils). **B**. CD11c^+^ CD11b^+^ CX3CR1^+^ (resident macrophages). **C**. CD11b^+^ Ly6C^+^ (inflammatory monocytes). **D**. CD11c^+^ CD103^+^ CD11b^-^ (left) and CD11c^+^ CD103^+^ CD11b^+^(middle), CD11c^+^ CD103^+^ (right) (dendritic cells). **E**. Lin^-^ CD90.2^+^ NK1.1^-^ KLRG1^+^ (ILC2). **F**. Lin^-^ CD90.2^+^ NK1.1^+^ KLRG1^-^ (ILC1). **G**. Lin-CD90.2^+^ KLRG1^-^ NK1.1^-^ CCR6^+^ (CCR6^+^ ILC3). **H**. Lin^-^ CD90.2^+^ KLRG1^-^ NK1.1^-^ RORγt^+^ (ILC3). **I**. Lin^-^ CD90.2^+^ KLRG1^-^ NK1.1^-^ RORγt^+^ IL-22^+^ (IL-22^+^ ILC3). **J**. Percentage of RORγt^+^ ILC3 (from G) expressing IL-22^+^. Lineage markers: CD3, CD5, B220, CD19, TCRβ, TCRγ8. Each symbol represents an individual mouse and horizontal bars indicate the median. Data combined from at least three independent experiments. Statistical analyses performed by Mann-Whitney. ns = not significant, \**p*≤0.05, \*\**p*≤0.01.

We have recently shown that *Hoil1^-/-^* mice have increased numbers of ILC2 in the small intestine (45), so we wondered if the impaired immune response to *C. rodentium* infection could be the result of altered ILC populations in the colon. The proportion of ILC2 was also increased in the colon of naïve *Hoil1^-/-^* mice, whereas proportions of NK1.1^+^ KLRG1^-^ ILC1 and CCR6^+^ ILC3 cells were reduced (Fig. 6E-G, Fig. S2E-F). Consistently, the type 2 inflammatory cytokine, *Il13*, was elevated in colon tissue from naïve *Hoil1*^-/-^ mice and during infection (Fig S3A). Since IL-13 can induce significant changes to the cellular composition and function of the intestinal epithelium (55), we wondered whether IL-13-dependent changes to the intestinal epithelium might contribute to the susceptibility of *Hoil1*^-/-^ mice to *C. rodentium*. To test this hypothesis, we used *Hoil1*^-/-^ mice crossed to *Il4ra^ff^* and Vil1-cre transgenic mice to block IL-13 signaling through the IL-4Rα selectively on intestinal epithelial cells (45). However, both *Hoil1^-/-^Il4ra^ff^* and *Hoil1^-/-^Il4ra^ΔIEC^* mice lost weight starting around 6 dpi, and succumbed to the infection between 10 and 15 dpi, whereas both *Hoil1^+/+^Il4ra^ΔIEC^* and *Hoil1^+/+^Il4ra^ff^*survived with infection to at least 21 dpi with minimal weight loss (Fig. S3B,C). Furthermore, *Hoil1^-/-^Il4ra^ΔIEC^* and *Hoil1^-/-^Il4ra^ff^* mice both shed more *C. rodentium* CFU in their stool at 2 dpi, and experienced increased systemic dissemination at 6 dpi relative to *Hoil1^+/+^Il4ra^ΔIEC^* and *Hoil1^+/+^Il4ra^ff^* littermates (Fig. S3D,E). These data indicate that the alterations in IEC differentiation caused by elevated IL-13 expression by ILC2 are not responsible for the increased replication, shedding and dissemination of *C. rodentium,* or morbidity observed in *Hoil1^-/-^* mice.

Further analysis of the colonic ILC3 populations indicated that naïve *Hoil1^-/-^* mice had fewer RORγt^+^ ILC3 and IL-22^+^ RORγt^+^ ILC3. IL-22^+^ RORγt^+^ ILC3 were also reduced in *Hoil1^-/-^* colon at 6 dpi (Fig. 6H,I, Fig. S2G). Interestingly, the percentage of RORγt^+^ ILC3 producing IL-22 did not appear to be significantly affected by HOIL1-deficiency before or during infection (Fig. 6J). Together, these data indicate that HOIL1 is important to regulate the number of ILC3 in the colon, and that a reduction in ILC3 numbers in the intestine of *Hoil1^-/-^* mice may contribute to defects in IL-22 production during *C. rodentium* infection, resulting in increased morbidity and mortality.

## Discussion

*C. rodentium* infection of mice is an excellent model system to interrogate the immune response to attaching and effacing bacterial gastrointestinal pathogens (7, 12). We have previously shown that HOIL1 is essential to prevent severe weight loss and lethality from *C. rodentium* infection, which causes colitis and intestinal hyperplasia with minimal morbidity in wild-type mice (24). In this study, we found that HOIL1 is required to restrict early bacterial replication, dissemination to systemic organs and damage to the colonic epithelium. The increased susceptibility of HOIL1-mutant mice was associated with impaired induction of IL-22 and a reduction in the proportion of colonic ILC3 before and during infection, indicating that HOIL1 is an important regulator of the type 3 inflammatory response.

HOIL1-deficient patients experience recurrent colitis and increased susceptibility to bacterial infections, including *E. coli* (40). HOIL1 is a component of the LUBAC. The other subunits are HOIP and SHARPIN, and loss of one subunit results in significantly impaired expression of the other two subunits. HOIP-deficient patients therefore experience similar symptoms (41, 42). HOIL1 is an E3 ubiquitin ligase, and its activity regulates HOIP activity, and limits the accumulation of unbranched glucosaccharides inside cells (38, 39). However, other substrates and functions of HOIL1 have been poorly defined to date. HOIP is also an E3 ubiquitin ligase, and generates methionine 1-linked (linear) polyubiquitin chains that facilitate NFκB activation downstream of multiple receptors including TLRs, TNFR and IL-1R (25-29, 32-34, 56). In mice, complete loss of HOIL1 or HOIP, or loss of HOIP E3 ligase activity is embryonic lethal, whereas mice expressing ligase-dead HOIL1 are viable (27, 37, 57, 58). In this study, we utilized two viable HOIL1-mutant mouse strains: *Hoil1*^-/-^ mice, that lack exons 8 and 9 encoding the RING1 domain (26); and *Rbck1*^f/-^ and *Rbck1*^-/-^ mice, in which exons 5 and 6 are flanked by loxP sites or removed by Cre-recombinase, where splicing from exon 4 to 7 results in a frameshift. Both *Hoil1*^-/-^ and *Rbck1*^-/-^ mice were highly susceptible to *C. rodentium* infection, and exhibited an increase in shedding in stool, infection of the distal colon and systemic dissemination within the first 4 days of infection. Importantly, *Rbck1^-/-^* mice were generated on a C57BL/6 background, ruling out contributions from susceptibility loci from other mouse strains that could be retained after backcrossing.

Although T cells and B cells are essential for the production of anti-*C. rodentium* antibodies for bacterial clearance and survival from infection (23, 59), *Hoil1*^-/-^ and *Rag1*^-/-^ *Hoil1*^-/-^ succumbed to infection significantly earlier than *Rag1*^-/-^*Hoil1*^+/+^ mice. Additionally, *Rag1*^-/-^*Hoil1*^-/-^ exhibited increased *C. rodentium* shedding in the stool and dissemination to systemic organs, demonstrating that HOIL1 plays a critical role in the innate immune response to infection. Although these findings do not rule out a role for HOIL1 in the adaptive response, we chose to focus our subsequent analyses on the innate response. Analysis of reciprocal bone marrow chimeric mice indicated that HOIL1 plays important roles in both radiation-resistant and radiation sensitive cells, but is more critical in radiation-resistant cells. Radiation-resistant cells include IECs and stromal cells, but also some populations of resident immune cells, such as ILCs. We were therefore surprised to find that loss of HOIL1 in IECs had no impact on the outcome of infection, especially since IEC are the main cell type exposed to and infected by *C. rodentium*. A recent study showed that HOIP activity is critical in IECs to prevent massive TNFα-mediated IEC death after injection with LPS (60). However, we observed very little induction of TNFα in the colon from our mice during infection, which might explain the lack of intestinal damage and weight loss in these mice. The same study also reported that mice lacking HOIP activity in IECs are not more susceptible to DSS-induced colitis, suggesting that physiological levels of TNFα in the colon during colitis may not be sufficient to induce wide-spread IEC death and loss of barrier integrity.

Instead, we determined that HOIL1 is required in lysozyme- and CD11c-expressing myeloid cells to limit bacterial dissemination and morbidity. Our findings are consistent with those of others that have demonstrated the importance of different subsets of DCs, monocytes, macrophages and neutrophils in limiting *C. rodentium*-induced weight loss and systemic dissemination (14, 50–52, 61–63). In the colon, CD11c-expressing cells include CD103^+^CD11b^-^ DCs, CD103^+^CD11b^+^ DCs and CD11b^+^CX3CR1^+^ resident macrophages, as well as infiltrating Ly6C^+^ monocytes and their derivatives during infection. CD103^+^CD11b^-^ DCs can produce IL-12 and support IFN-γ-mediated protection of the mucosal barrier (53). Despite the reduction in CD103^+^CD11b^-^ DCs in the colon of *Hoil1*^-/-^ mice, we were unable to detect differences in IFN-γ protein or mRNA expression at 6 dpi, suggesting that the IL-12 – IFN-γ pathway was not significantly altered. CD103^+^CD11b^+^ DCs, located primarily in isolated lymphoid follicles, produce IL-23 and support the production of IL-22 by ILC3 and Th22 cells (14, 51, 53). Although we did not observe a reduction in CD103^+^CD11b^+^ DCs or IL-23 expression, it is possible that loss of HOIL1 in these cells impairs their expression of cytokines and other molecules that support the maintenance and functions of ILC3. CD11b^+^CX3CR1^+^ resident macrophages play an important role in maintenance of the mucosal barrier. They are highly phagocytic and bactericidal against bacteria that breach the epithelial barrier, are able to sample the intestinal lumen with transepithelial dendrites, yet are hyporesponsive to TLR agonists and maintain an anti-inflammatory phenotype (54, 64–66). Resident macrophages also constitutively produce IL-1β that supports intestinal homeostasis (67). Although we did not observe any differences in the numbers of CD11b^+^CX3CR1^+^ cells or expression of IL-1β in the colon, it is possible that their bactericidal activity is impaired in the absence of HOIL1. Macrophages derived from infiltrating monocytes are more pro-inflammatory. However, we did not observe major differences in the expression of cytokines in the colon at 6 dpi, except for reductions in IL-22 and CCL20, and an increase in IL-6. Lysozyme M-expressing cells in the colon include neutrophils and macrophages, which are important early responders to *C. rodentium* infection due to their bactericidal activity and production of inflammatory mediators (61, 62). A possible role for HOIL1 in bactericidal activity is supported by the lack of detectable increase in intestinal permeability at 4 dpi, at which time *C. rodentium* CFU were detected in the spleen and liver.

The loss of HOIL1 in myeloid cells does not fully account for the susceptibility of *Rbck1*^-/-^ mice, or of the WT->KO bone marrow chimeric mice, indicating that another cell type is involved. It is likely that HOIL1 plays important roles in multiple cell types during *C. rodentium* infection. HOIL1-deficiency in IECs may have a greater impact on intestinal pathology when HOIL1 is also absent in other cell types that support the IEC barrier function, such as macrophages or ILC3. It will be interesting to determine the requirement for HOIL1 expression in ILC3 for their maintenance and function.

Of note, we observed a significant decrease in the expression of CCL20, a chemokine produced constitutively by IECs and induced in macrophages and IECs. CCL20 can be induced by pro-inflammatory cytokines, including IL-1β, and recruits CCR6-expressing cells, which include CCR6^+^ ILC3, LTi cells, Tγ8 cells and immature DCs (68, 69). Indeed, we observed a reduction in the number of CCR6^+^ and RORγt^+^ ILC3 in the colon of *Hoil1*^-/-^ mice before and during infection. This was accompanied by an increase in ILC2, and therefore may reflect a shift in the recruitment, differentiation or plasticity of ILCs in the colon. Production of IL-22 by ILC3 is critical for maintenance of barrier integrity and survival from *C. rodentium* infection (17, 70), and we observed a 50% reduction in IL-22 protein during infection, and 2-3-fold fewer IL-22^+^ ILC3 in the colon before and during infection that may be at least partially responsible for the increased susceptibility of *Hoil1*^-/-^ mice. The increase in ILC2 numbers was associated with elevated IL-13 in the colon, which we had previously observed in the small intestine (45). IL-13 signaling in IECs induces changes to IEC proliferation and differentiation, including an increase in goblet cells and Paneth cells in the small intestine, and alters the production of mucins and anti-microbial peptides. These changes to the mucosal and luminal environment may impact nutrient availability, commensal microbes, as well as colonization with pathogens. We wondered if these IL-13-dependent changes might be responsible for the increase in early colonization, replication and shedding of *C. rodentium* in *Hoil1*^-/-^ mice. However, deletion of the IL-4Rα subunit of the IL-13 receptor in IECs did not significantly ameliorate early shedding, dissemination, weight loss or overall mortality of *Hoil1*^-/-^ mice, indicating that IL-13-dependent changes to IEC differentiation and function do not play a significant role. However, this does not rule out a role for type 2 inflammatory cytokines in modulating the function of other cell types, such as macrophages.

Additional studies will be required to determine whether HOIL1 or HOIP E3 ligase activity is required for protection during infection. Of note, MyD88-deficient mice are similarly susceptible to *C. rodentium*, suggesting that the generation of linear ubiquitin chains by HOIP is important (61, 71–74). MyD88 functions downstream of most TLRs as well as the IL-1 receptor family that includes IL-1R, IL-18R, IL-33R (ST2), and IL-36R. TLR2-deficiency is also detrimental during *C. rodentium* infection, although TLR4-deficiency appears to be protective by limiting intestinal inflammation (75–77). The IL-1R is also important, particularly on ILC3 and on IECs for the induction of IL-22 and response to IL-22, respectively, as well as the induction of chemokines, such as CCL20 (15, 16, 68, 78). IL-36R signaling also promotes IL-22 induction during *C. rodentium* infection through enhancement of IL-23 and IL-6 expression (79). However, the phenotypes of *Myd88*^-/-^ and *Hoil1*^-/-^ mice are not identical: hyperplasia is almost completely absent in *Myd88*^-/-^ mice, but apparent in *Hoil1*^-/-^ mice; and we observed that HOIL1 was more important in radiation-resistant cells, whereas MyD88 is more important in bone marrow-derived cells (61, 71). These findings indicate that HOIL1 and LUBAC activity may be required in additional signaling pathways.

Overall, we have found that HOIL1 and the LUBAC play an important role in promoting a type 3 inflammatory response during intestinal infection with a bacterial pathogen. Type 3 responses are critical for controlling other bacterial pathogens, such as EPEC and *Salmonella*, as well as for modulating responses to the commensal microbiota. The regulation of ILC3 numbers in naïve mice suggests that these findings may also be relevant to IBD, where translocation of commensal microbes may be increased (80). A more thorough understanding of these mechanisms may aid in the development of therapeutics for gastrointestinal bacterial infections and inflammatory disorders.

## Funding Information

This work was funded in part by UIC institutional start-up funds and by NIAID R01 AI150640 to D.A.M.

## Acknowledgements

We would like to acknowledge D. Kreamalmeyer, M. White, X. Zhang, J. Marshall, M. Byrne, S. Madathil (Millipore), the UIC Research Resources Center Histology and Tissue Imaging Core and the Flow Cytometry Core, and the Digestive Diseases Research Core Center and the Developmental Biology Histology Core at Washington University for technical assistance. We thank members of the MacDuff laboratory and Skip Virgin’s laboratory for helpful discussions, and Bruce Vallance for helpful suggestions. We thank Center d’ImmunoPhenomique (Ciphe) for providing the mutant mouse line (Allele: Rbck1^tm1a(EUCOMM)Hmgu^), INFRAFRONTIER/EMMA (www.infrafrontier.eu, PMID: 25414328), and Institut de Transgenose (INTRAGENE, Orleans, France) from which the mouse line was distributed (EM:09852) (46, 47). Associated primary phenotypic information may be found at www.mousephenotype.org.

---

**Supplementary Figure 1: An independent *Hoil1* (*Rbck1*) mutant mouse strain exhibits similar susceptibility to *C. rodentium*. A**. Western blots for the indicated LUBAC components and actin on spleen lysates from *Rbck1*^f/f^ and *Rbck1*^-/-^ mice. Each lane contains a sample from an independent mouse. **B**. Relative *Hoil1/Rbck1* mRNA levels in distal colon (left) and spleen (right) detected with primer-probe sets within the indicated exons. **C-D**. Percent initial body weight (C) and survival (D) of *Rbck1*^f/f^ (circles, n=10) and *Rbck1*^-/-^ (squares, n=7) mice following i.g. inoculation with 2x10^9^ CFU *C. rodentium.* **E-G**. *C. rodentium* CFU in stool (E), distal colon (F), liver, spleen, and MLN (G) at the indicated times post-infection. **H**. Relative *Hoil1/Rbck1* mRNA levels in spleen and colonic IECs from *Rbck1*^f/f^ and *Rbck1*^f/f^ ^Vil1cre^ mice detected with primer-probe sets within the indicated exons. For B and E-H, each symbol represents an individual mouse, and the median is indicated. Dotted lines indicate the LOD. Data were combined from at least three independent experiments. Statistical analyses were performed by log ranked Mantel Cox (D), multiple t-tests (C), or Mann-Whitney tests (B, E-H). \**p*≤0.05, \*\**p*≤0.01.

**Supplementary Figure 2: Gating strategies and representative flow cytometry plots related to** **Figure 5**. Representative flow cytometry plots for colonic lamina propria cells from *Hoil1^+/+^* (left) and *Hoil1^-/-^* (right) mice at baseline (top row) or 6 dpi with *C. rodentium* (bottom row). **A**. Pre-gating on single, live, CD45+ cells. **B**. CD11b and Ly6G expression**. C**. CD11b and CX3CR1 expression, pre-gated CD11c^+^ cells (right plot). **D**. CD11b expression, pre-gated on CD11c^+^, CD103^+^ cells (right plot). **E**. KLRG1 and NK1.1 expression, pre-gated on lineage^-^, CD90.2^+^ cells (upper plot). **F.** CCR6 expression, gated on lineage^-^, CD90.2^+^, KLRG1^-^, NK1.1^-^ cells (E). **G**. RORγt and IL-22 expression, pre-gated on CD90.2^+^, KLRG1^-^, NK1.1^-^ cells (E). Flow plots are representative data from at least three independent experiments. Lineage markers: CD3, CD5, B220, CD19, TCRβ, TCRγ8.

**Supplementary Figure 3: Blocking IL-13 signaling via IL-4Rα in IECs does not protect *Hoil1*^-/-^ mice from *C. rodentium* pathogenesis. A.** *Il13* mRNA expression in the distal colon of naïve *Hoil1*^+/+^ and *Hoil1*^-/-^ mice. B-C. Percent initial body weight (B) and survival (C) of *Hoil1^-/-^Il4ra^ff^* (*n*=10), *Hoil1^-/-^Il4ra^ΔIEC^* (*n*=4), *Hoil1^+/+^Il4ra^ff^* (*n*=6) and *Hoil1^+/+^Il4ra^ΔIEC^* (*n*=13) mice at the indicated days post-infection. D-E. *C. rodentium* CFU in stool at 2 dpi (D) and spleen, liver and MLN from *Hoil1^-/-^Il4ra^ff^*, *Hoil1^-/-^Il4ra^ΔIEC^*, *Hoil1^+/+^Il4ra^ff^* and *Hoil1^+/+^Il4ra^ΔIEC^* mice at 6 dpi. Median values are indicated. Dashed lines indicate the limit of detection. Statistical analyses were performed by Mann Whitney test (A) or Kruskal-Wallis test with Dunn’s multiple comparison test (D-E). \**p*≤0.05, \*\**p*≤0.01, \*\*\**p*≤0.001.

## Notes

### Competing Interest Statement

The authors have declared no competing interest.

