## Supplementary figures and images for "HOIL1 regulates group 3 innate lymphoid cell numbers in the colon and protects against systemic dissemination, colonic ulceration, and lethality from *Citrobacter rodentium* infection"

### Supplemental Figure 1

#### Supplemental Figure 1

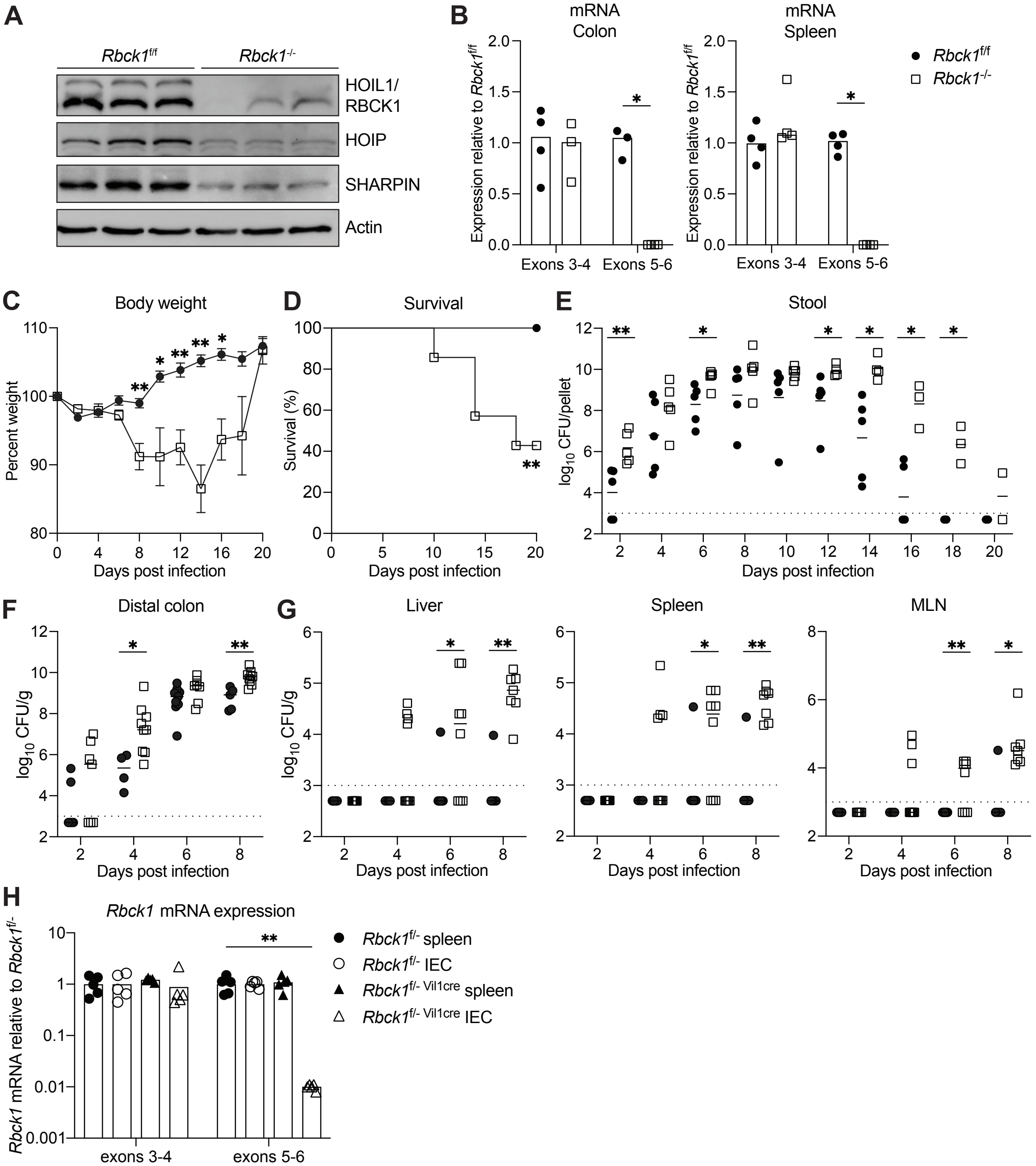

### Supplemental Figure 2

#### Supplemental Figure 2

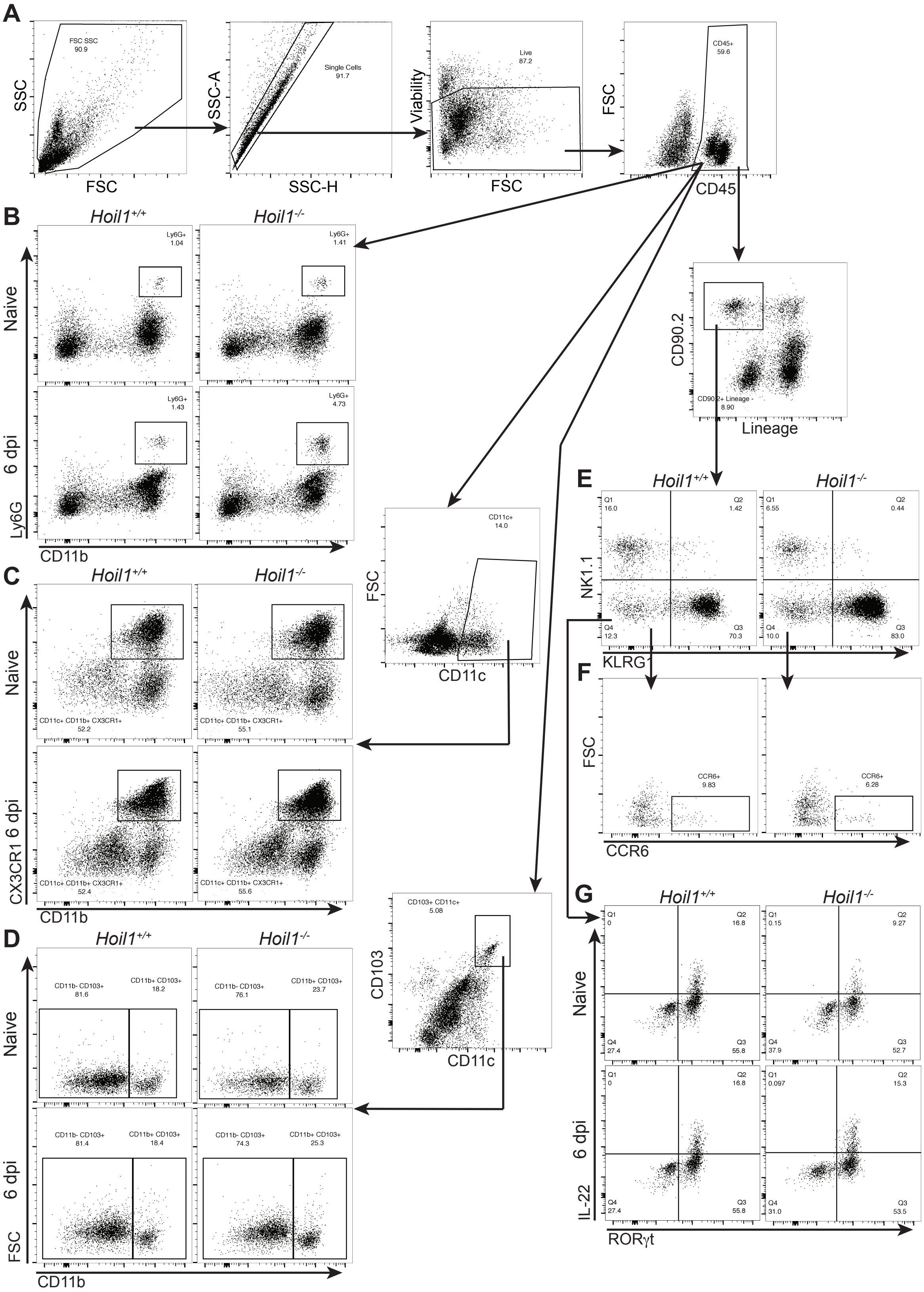

### Supplemental Figure 3

#### Supplemental Figure 3

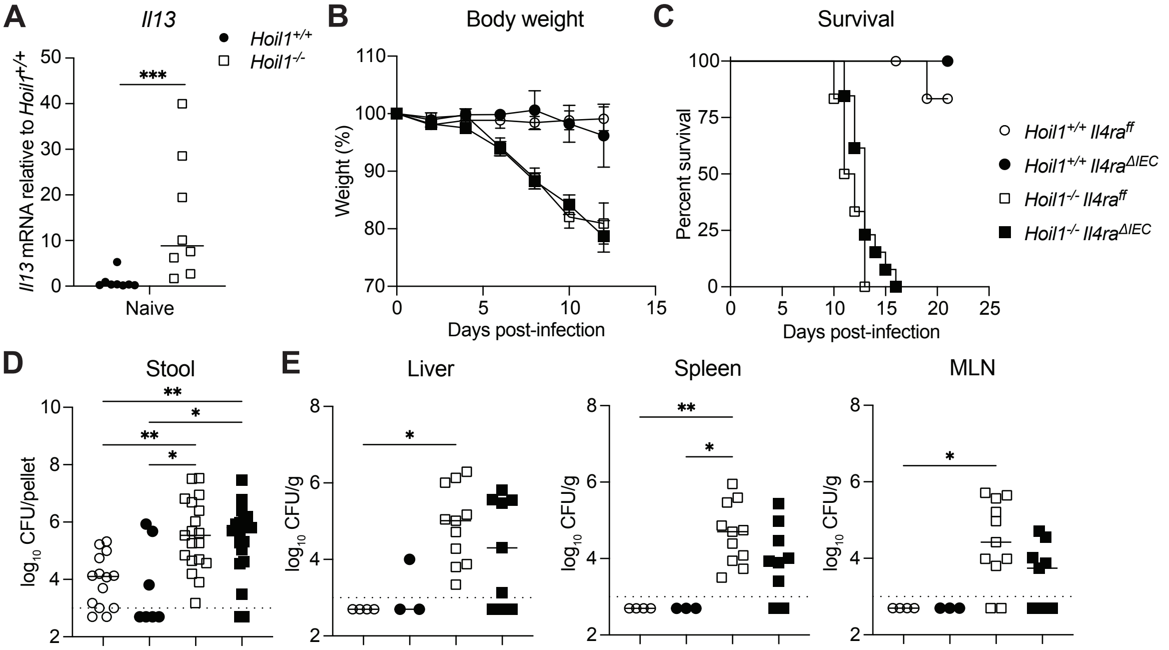
